# Bayesian Encoding and Decoding as Distinct Perspectives on Neural Coding

**DOI:** 10.1101/2020.10.14.339770

**Authors:** Richard D. Lange, Sabyasachi Shivkumar, Ankani Chattoraj, Ralf M. Haefner

## Abstract

One of the most influential, and controversial, ideas in neuroscience has been to understand the brain in terms of Bayesian computations. Unstated differences in how Bayesian ideas are operationalized across different models make it difficult to ascertain both which empirical data support which models, and how Bayesian computations might be implemented by neural circuits. In this paper, we make one such difference explicit by identifying two distinct philosophies that underlie existing neural models of Bayesian inference: one in which the brain recovers experimenter-defined structures in the world from sensory neural activity (Decoding), and another in which the brain represents latent quantities in an internal model that explains its inputs (Encoding). These philosophies require profoundly different assumptions about the nature of inference in the brain, and lead to different interpretations of empirical data. Here, we characterize and contrast both philosophies in terms of motivations, empirical support, and relationship to neural data. We also show that this implicit difference in philosophy underlies some of the debate on whether neural activity is better described as a sampling-based, or a parametric, distributional code. Using a simple model of primary visual cortex as an example, we show mathematically that it is possible that the very same neural activity can be described as probabilistic inference by neural sampling in the *Encoding* framework while also forming a linear probabilistic population code (PPC) in the *Decoding* framework. This demonstrates that certain families of Encoding and Decoding models are compatible with each other rather than competing explanations of data. In sum, Bayesian Encoding and Bayesian Decoding are distinct, non-exclusive philosophies, and appreciating their similarities and differences will help organize future work and allow for stronger empirical tests about the nature of inference in the brain.

## 1 Introduction

According to the Bayesian Brain hypothesis, neural circuits carry out statistical computations by combining prior knowledge with new evidence, combining multiple sources of information according to their reliability, and taking actions that account for uncertainty. In the case of perception, prior knowledge is assumed either to come from experience with the world during development or to be encoded genetically, having been learned over the course of evolution. While any given sensory measurement may be noisy or ambiguous – providing a wide likelihood function in Bayesian terms – prior knowledge is deployed to resolve these ambiguities when possible (von Helmholtz, 1925). The Bayesian framework has been instrumental for our understanding of perception and perceptual decision-making (Knill and Richards, 1996; Kersten et al., 2004; Fiser et al., 2010; Pouget et al., 2013).

At the core of the Bayesian Brain hypothesis is the idea that neural activity corresponds to probability distributions rather than point estimates – such schemes are known as “distributional codes” (Zemel et al., 1998). Previous surveys of distributional codes have emphasized a distinction between sampling-based and parametric codes (Fiser et al., 2010; Pouget et al., 2013; Sanborn, 2015; Gershman and Beck, 2016). From a computational standpoint, sampling and parametric codes each have advantages and disadvantages. In the context of neuroscience, sampling and parametric codes have also been compared with respect to the simplicity of implementing computations believed to be important for the brain, such as cue combination and marginalization (Fiser et al., 2010). Further, numerous studies have empirically tested for properties of sampling or parametric codes in neural responses. Sampling codes have been used to explain spontaneous cortical activity (Berkes et al., 2011), neural variability (Hoyer and Hyvärinen, 2003; Orban et al., 2016; Festa et al., 2021), structure in noise correlations (Haefner et al., 2016; Bányai et al., 2019), and onset transients and oscillations (Aitchison and Lengyel, 2016; Hennequin et al., 2018; Echeveste et al., 2020). Meanwhile, parametric codes have been cited in explanations of contrast-invariant tuning curves (Ma et al., 2006), near-linearity during cue-combination (Fetsch et al., 2011, 2013), evidence integration dynamics in parietal cortex (Beck et al., 2008; Hou et al., 2019), divisive normalization (Beck et al., 2011), and more (Pouget et al., 2013). Importantly, sampling and parametric codes have so far always been discussed and compared as competing and mutually exclusive mathematical models of the same neural circuits, with no decisive evidence presented favoring one over the other model. Notably, contrast-invariant tuning and divisive normalization have also been replicated by sampling models (Orbán et al., 2016; Echeveste et al., 2020).

The primary goal of this paper is to characterize and contrast two distinct perspective on the Bayesian Brain hypothesis, which we call **Bayesian Encoding** and **Bayesian Decoding**. These are complementary perspectives that make different assumptions about the nature of the inference problems faced by the brain, and are supported or falsified by different kinds of empirical data. We argue that not making their differences explicit has led to confusion about how to interpret empirical data. In particular, we describe how the above debate on whether neural responses are better modeled as samples or parameters is complicated by the fact that sampling codes usually make assumptions consistent with Bayesian Encoding while parametric codes often make assumptions consistent with Bayesian Decoding. However, neither the connection between Bayesian Encoding and sampling, nor between Bayesian Decoding and parametric codes, is a necessary consequence of either theory. Indeed, there are Encoding models built on parametric codes, Decoding models based on sampling, and still other models that contain elements of both approaches (Section 2.4 and Table 1 below).

**Table 1:**
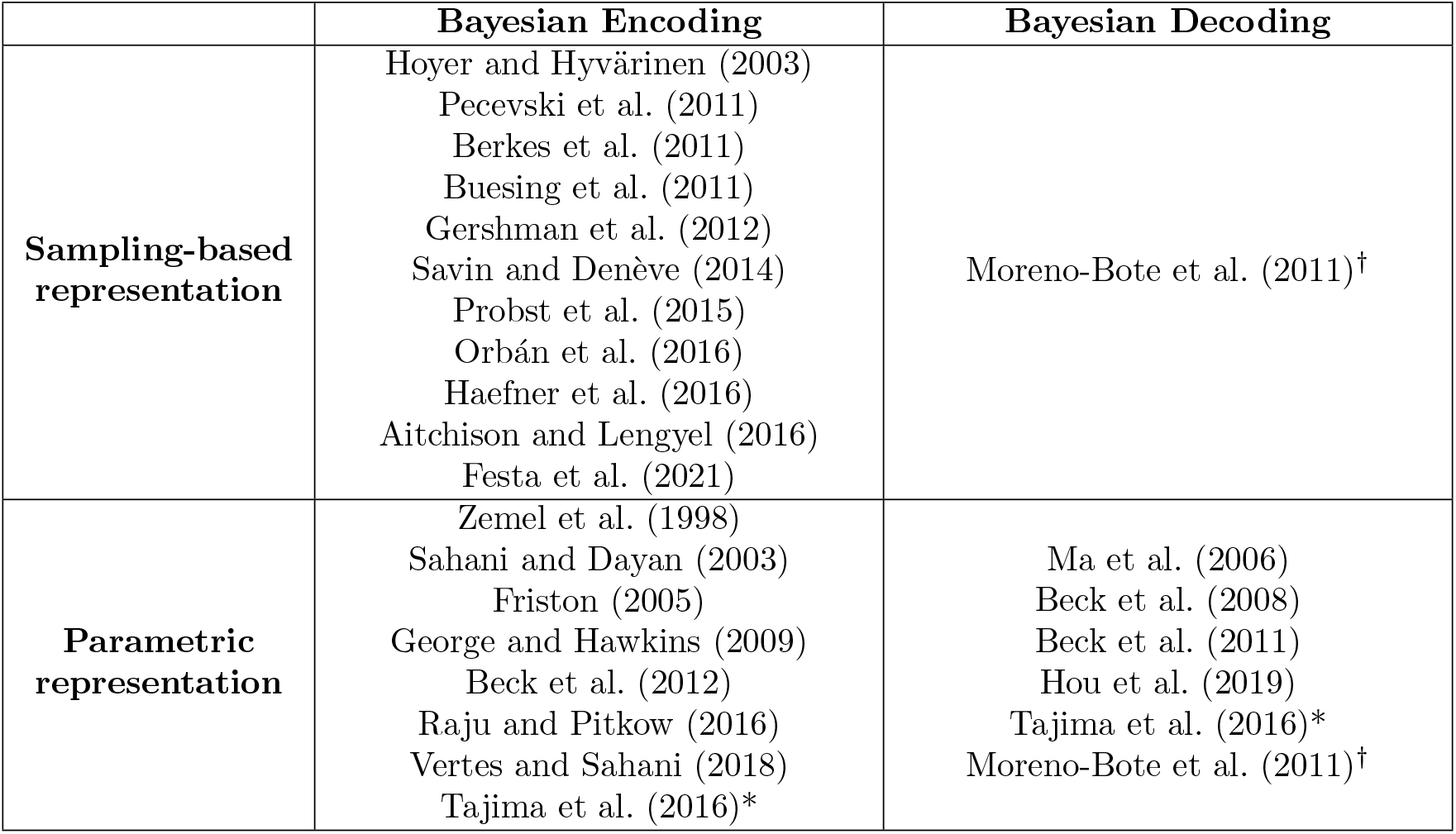
Classifying previous work on Bayesian neural models according to whether they construct Bayesian Encoding or Decoding models, and whether they use a sampling-based or a parametric neural representation. Tajima et al., marked with “*” contains elements of both encoding and decoding. Moreno-Bote et al., marked with “†”, contains elements of both sampling-based and parametric decoding.

Finally, we illustrate the complementary nature of these two philosophies using a simple model of primary visual cortex (Shivkumar et al., 2018). In this example, we construct a sampling-based *Encoding* model based on a linear Gaussian model of natural images (Olshausen and Field, 1996, 1997), and derive the implied *Decoding* model. We show that firing rates in this model form a canonical kind of parametric code: a Probabilistic Population Code (PPC). There is thus no inherent contradiction in saying that the brain is *both* sampling (in the “Bayesian Encoding” sense) and represents parameters (in the “Bayesian Decoding” sense), and depending on the encoding models’ generative model, and the considered task, this parametric code may even be a linear PPC. We conclude with a discussion of distributional neural codes in general.

## 2 Bayesian Encoding vs Bayesian Decoding

We follow the seminal work of Zemel et al. (1998) in assuming that patterns of neural activity represent entire probability distributions over a variable, not just a point estimate of it, i.e. that they form a *distributional* code. The nature of this “variable” and its relationship to neural response is key to the distinction between the Bayesian Encoding and the Bayesian Decoding frameworks.

### 2.1 Bayesian Encoding

We define **Bayesian Encoding** as the view that there exists a probability distribution over some quantity of potential interest to the brain, and that the primary function of sensory neurons is to compute and represent an approximation to this distribution. We use the term “encoding” because the probability distribution that neurons are hypothesized to represent conceptually precedes the actual neural responses. That is, in Bayesian encoding models, there exists a *reference distribution* that is defined independently of how neurons actually respond, and which is approximately encoded by neural responses.

Bayesian Encoding requires a source for the reference distribution. In the context of the sensory system, this typically takes the form of an internal generative model of sensory inputs, and the distribution to be encoded is the posterior over latent variables in that model (Figure 1a-b). With this perspective, the goal of sensory areas of the brain is to learn a statistical model of its sensory inputs (Dayan et al., 1995; Dayan and Abbott, 2001; Fiser et al., 2010; Berkes et al., 2011) in which sensory observations, such as an image on the retina, are explained as the result of higher order causes. Whereas the information on the retina is highly mixed – objects, lights, textures, and optics interact in complex ways to create an image – the internal model aims to explain sensory data in terms of unobserved causes that are often assumed to be sparse and independent (von Helmholtz, 1925; Olshausen and Field, 1996; Bell and Sejnowski, 1997). A generative model makes this process explicit by assigning prior probabilities to the (co)occurrence of causes (represented by latent variables) and by quantifying the likelihood of a particular configuration of the causes for generating a particular sensory observation. The encoded posterior distribution in this framework is defined over the latent variables in this statistical model.

**Figure 1:**
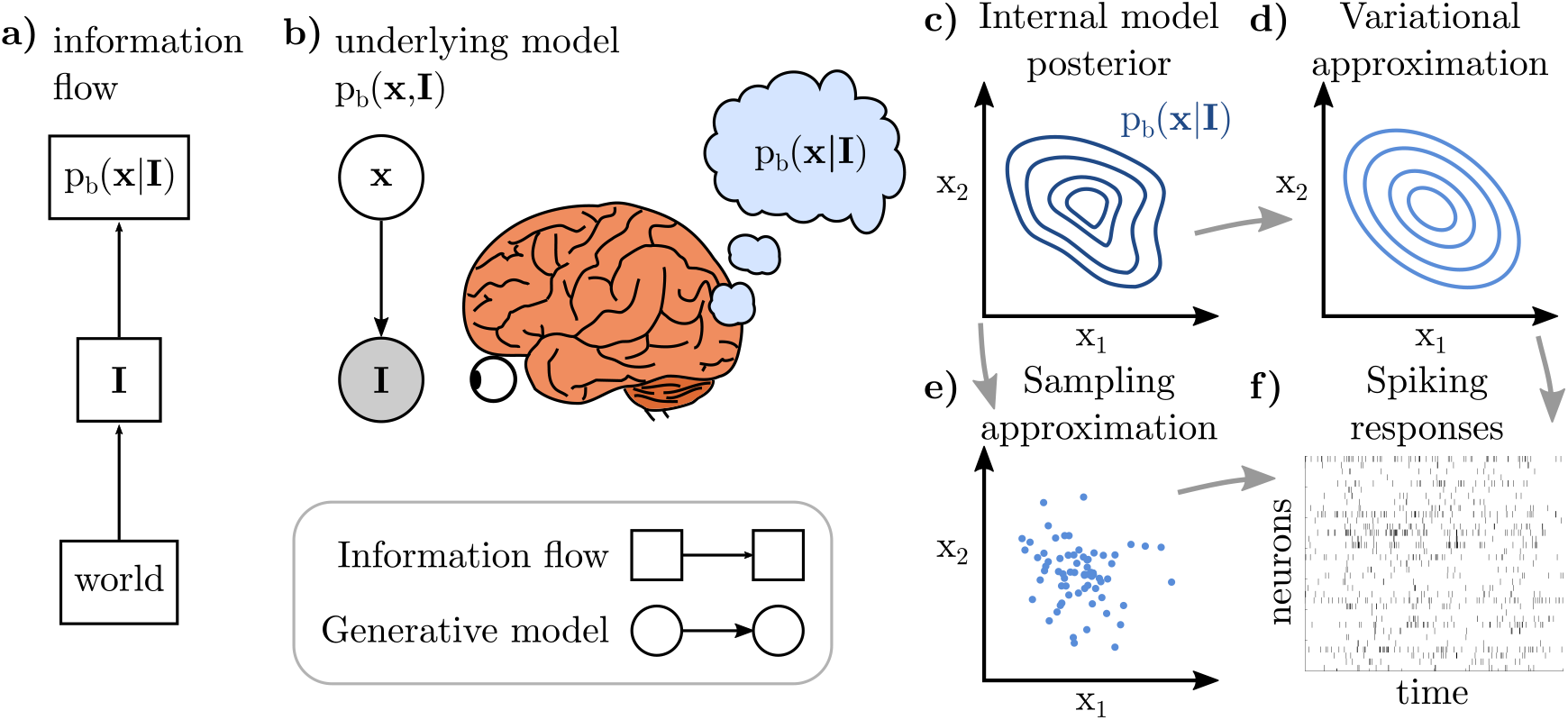
Visualization of Bayesian Encoding. **a)** Diagram of information flow: the world provides sensory inputs (**I**), which then give rise to inferences about latent variables (**x**). **b)** Bayesian Encoding typically assumes that the brain has an internal model of its inputs, and that perceptual inferences are about variables in this internal model, not necessarily corresponding to quantities in the external world *per se.* With Bayesian Encoding, it is also typical to assume that the internal model is *task-independent* and that the brain always computes a posterior over internal variables, p_b_ (**x**|**I**), regardless of whether **I** is a highly controlled stimulus in a task or encountered in the wild. **c-f)** The defining feature of Bayesian Encoding is the existence of a reference distribution (c), typically the posterior over a set of latent variables, **x**, given a sensory measurement, **I**. One then assumes an approximation scheme such as variational inference (c→d) or sampling (c→e), and that this approximation is then realized in patterns of neural activity (f).

For latent variables **x** and sensory input **I**, optimal inference means computing the posterior distribution,

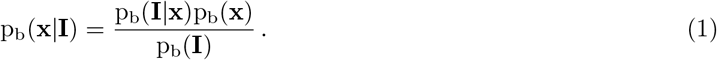

We use the subscript b in p_b_(·) to refer to quantities in the brain’s internal model, and to distinguish them from other types of probabilities such as a decoder’s uncertainty. A prototypical case of Bayesian Encoding poses the question of how neural circuits could compute and represent the posterior distribution p_b_(**x**|**I**) for any sensory **I**, given the internal model that the brain has learned (Figure 1c), and how it can learn this internal model in the first place. In general, exact inference is an intractable problem (Murphy, 2012; Wainwright and Jordan, 2008; Bishop, 2006), leading to the question of how the brain could compute and represent an *approximation* to the true posterior (Figure 1d-f), and what the nature of this approximation is. This line of reasoning motivates work on “neurally plausible approximate inference algorithms,” including approaches with connections to sampling-based inference (Figure 1e), as well as approaches inspired by variational inference techniques, related to parametric neural codes (Figure 1d) (reviewed in Fiser et al. (2010); Sanborn (2015); Gershman and Beck (2016)).

### 2.2 Bayesian Decoding

We define **Bayesian Decoding** as the perspective in which neural activity is treated as *given*, and emphasis is placed on the statistical uncertainty of a decoder observing those neural responses. Bayesian Decoding is closely related to ideal observer models in psychophysics, involving tasks that require the estimation of scalar aspects of a presented stimulus (e.g. its orientation or its contrast) or a decision whether the stimulus belongs to one of two or more discrete classes (e.g. “left” or “right”). Of course, any stimulus *s* that elicits neural responses **r** is optimally decoded by computing p(*s*|**r**) (Figure 2). The key question within the Bayesian Decoding framework is this: what conditions must the stimulus-driven neural activity (p(**r**|*s*)) fulfill such that the decoder (p(*s*|**r**)) is *simple,* e.g. linear and invariant to nuisance? For instance, imposing linearity and invariance constraints on the decoder implies constraints on tuning curves and the distribution of neural noise (Zemel et al., 1998; Ma et al., 2006).

**Figure 2:**
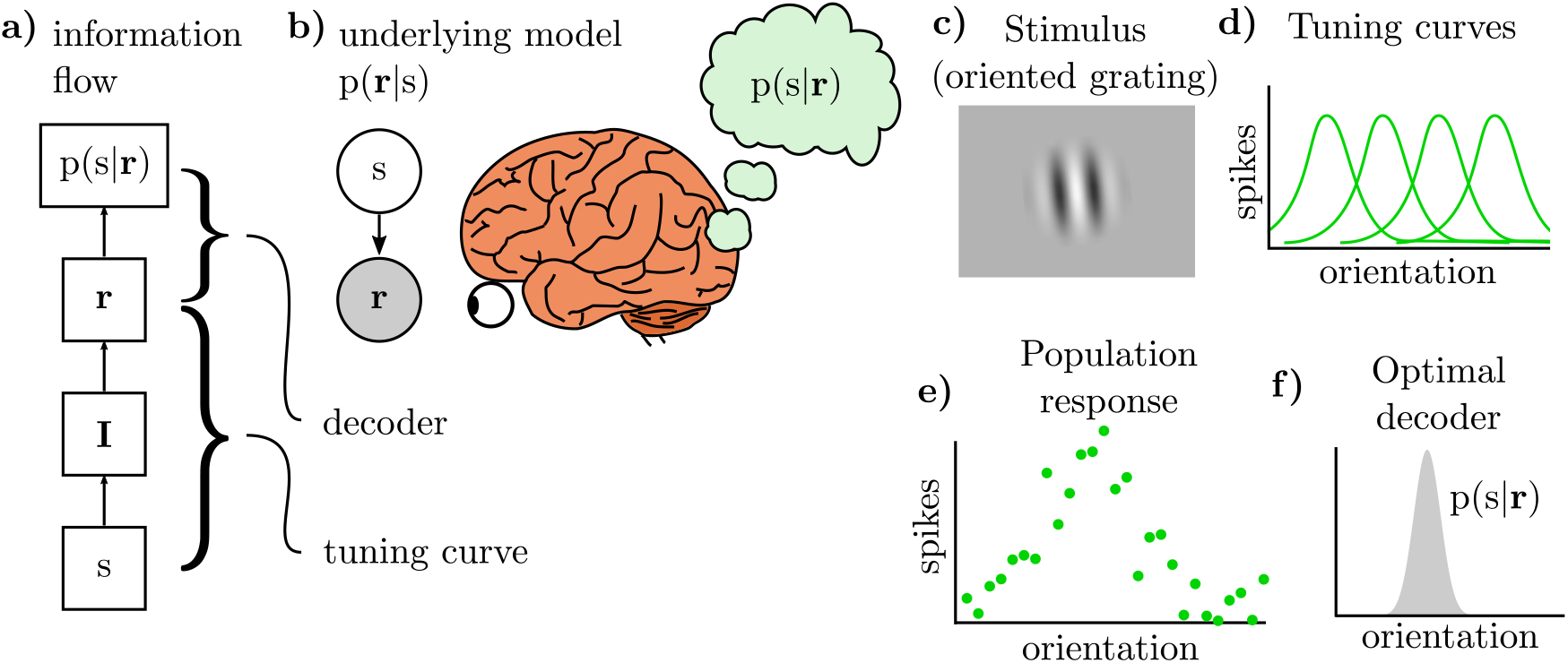
Visualization of Bayesian Decoding. **a)** Diagram of information flow: a quantity of interest (*s*) in the world elicits neural responses (**r**), mediated by sensory inputs (**I**). The decoding question is how the brain forms an internal estimate, *s*, from **r**. **b)** The underlying probabilistic model assumes that *s* generates **r**, so inference is the problem of recovering *s* from **r**. **c)** The decoding problem usually begins with a stimulus, such as the orientation of a grating of a given spatial frequency, size, location, and contrast. **d-f)** Given a population of neurons’ tuning curves to *s* (d) and an observation of spikes on a single trial (e), an optimal decoder computes p(*s*|**r**) (f).

Bayesian Decoding is closely related to familiar notions of optimal neural decoding. Classically, decoding is either a tool for assessing information content in neural responses or a mechanistic model of how they impact behavior. In the Bayesian setting, the emphasis is on how neural activity is interpreted by the rest of the brain and influences behavior, and how this depends on the brain’s uncertainty about a behaviorally-relevant stimulus.

Probabilistic Population Codes (PPCs), as introduced by Ma et al (2006), exemplify the Bayesian Decoding approach. PPCs construct a Bayesian decoder that is both simple and invariant to nuisance: if a population of neurons tuned to *s* has “Poisson-like” variability, then the optimal decoder is part of the exponential family with firing rates as natural parameters. This is a particularly convenient representation for taking products of two distributions as required by cue-integration (Ma et al., 2006) and evidence accumulation Beck et al. (2008). Equally important is the notion of *invariance* afforded by a PPC: as long as nuisance variables such as image contrast or dot coherence only multiplicatively scale tuning curves, the decoder can ignore them.

Importantly, under the assumption that the brain employs a computationally convenient neural code, linearity for cue combination and multiplicative gain by nuisance variables become *predictions* of PPCs. In classical decoding approaches, neural responses are simply “given,” not prescribed by a theory. In the Bayesian Decoding framework generally, and in the case of PPCs in particular, imposing constraints on the decoder constrains the possible set of evoked response distributions, p(**r**|*s*). These constraints have then been formulated as predictions and tested empirically (Fetsch et al., 2011, 2013; Pouget et al., 2013; Hou et al., 2019).

### 2.3 Contrasting Bayesian Encoding and Bayesian Decoding

There are four key differences between the Bayesian Encoding and Bayesian Decoding perspectives, which we will discuss in each of the following sections: (1) what they assume the brain is inferring, (2) what the terms “likelihood” and “posterior” refer to, (3) the role of neural responses in the theory, and (4) the empirical data and other arguments used to motivate them. As our goal is to summarize and categorize a large and diverse sub-field, there will be exceptions to each rule, but we expect these distinctions to be useful for framing further discussions.

#### 2.3.1 Differences in what is assumed to be inferred

An integral part of the Bayesian Encoding framework is the existence of an abstract internal model that could in principle be implemented *in silico* or in the brains of other individuals or other species. Deriving predictions for neural data requires an additional linking hypothesis on the nature of distributional codes, such as whether neurons sample or encode variational parameters, and how either samples or parameters correspond to observable biophysical quantities like membrane potentials and spike times or spike counts. Bayesian Encoding thus decomposes the question of what sensory neurons compute into two parts: first, what is the internal model which defines optimal inference (the reference distribution), and second, how do neural circuits carry out approximate inference in that model (e.g. sampling or parametric)?

The brain’s internal model is typically assumed to have been calibrated through exposure to natural stimuli (Dayan et al., 1995; Dayan and Abbott, 2001; Berkes et al., 2011) and to only change slowly with exposure to new stimuli in adult brains. For this reason, the generative model in Bayesian Encoding models, especially in the case of early sensory areas, is often assumed to be independent of experimental context. For instance, if the brain’s internal model comprises patches of local image features, then it is assumed that the brain infers and encodes the same set of image features, whether viewing natural scenes or artificial stimuli in a task (Haefner et al., 2016; Orbán et al., 2016; Shivkumar et al., 2018; Bányai et al., 2019). The assumption of calibration in a Bayesian Encoding framework also makes predictions for how the internal model should change in response to the statistics of sensory inputs during development (Berkes et al., 2011), and to extensive exposure to stimuli in a particular task (Lange and Haefner, 2022).

In contrast, Bayesian Decoding models are typically applied in the context of estimating task-relevant variables. For instance, in a motion discrimination task, a Bayesian Decoding question would be how the brain represents uncertainty over directions of motion. Importantly, answering this question does not require a generative model of all possible naturally-occurring motion stimuli, nor does it require a true or correct reference distribution over stimuli; it requires only a statistical model of the relation between scalar motion direction in a particular task (and possibly nuisance variables like coherence) and neural responses, p(**r**|*s*), making it observable experimentally. The difference between (typical realizations of) the Bayesian Encoding and Decoding perspectives is illustrated in Figure 3.

**Figure 3:**
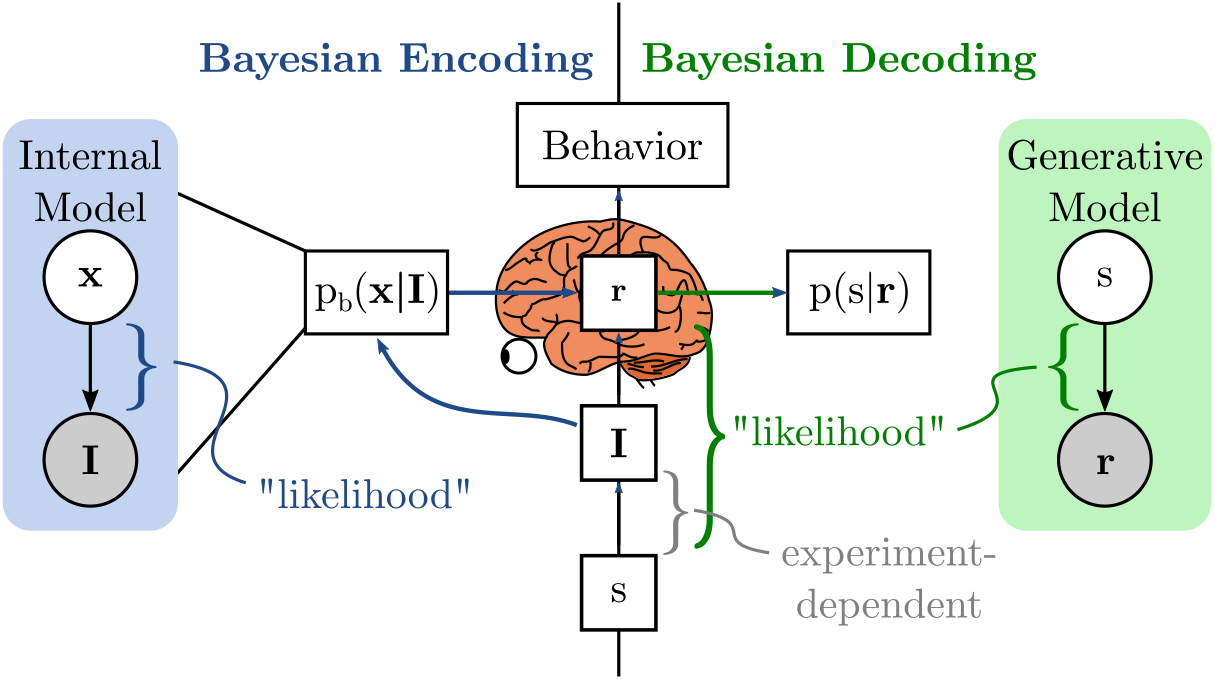
Side-by-side comparison of Bayesian Encoding and Bayesian Decoding. In both frameworks, it is understood that there exists a mechanistic, biophysical connection between stimuli (**I**), sensory neural responses (**r**), and behavior. In the Bayesian Decoding framework, emphasis is placed on the uncertainty of a decoder estimating a stimulus parameter *s* from **r** (green arrow). Bayesian Encoding posits the existence of an internal model with latent variables **x**, and that neural responses, **r**, encode the computation of a posterior distribution, p_b_(**x**|**I**). The blue arrow from p_b_(**x**|**I**) to **r** can be seen as an instance of *downward causation* between levels of abstraction, where changes to the posterior (at the algorithmic level) imply changes to neural responses (at the implementation level) (Campbell, 1974; Yablo, 1992; Lange and Haefner, 2022). In Bayesian Decoding, the “likelihood” refers to p(**r**|*s*), and the inference problem is to recover *s* from **r**. In Bayesian Encoding, the “likelihood” refers to the internal model’s p_b_(**I**|**x**), and the inference problem is to recover **x** from **I** and to embed the posterior over **x** in **r**. In any psychophysical task, the link between *s* and **I** depends on the experiment (gray bracket). Importantly, this means that the “likelihood” in a Bayesian Decoding model depends on choices made by the experimenter (such as their choice of stimuli), but not in a Bayesian Encoding model.

We emphasize that Bayesian Encoding *typically* but not *necessarily* involves a task-independent internal generative model, and Bayesian Decoding likewise is *typically* but not *necessarily* applied to task-specific variables. In fact, where the ideas of encoding and decoding distributions first appeared in Zemel et al (1998), the encoded distribution was over task quantities (such as **x** being motion or heading direction), without specifying an internal generative model, and the decoding problem was framed as the inverse to the encoding problem – that is, recovering the encoded p(**x**| …) from **r**. This again emphasizes the complementary nature of these philosophies: we are free to apply the Bayesian Decoding framework to variables in a task-independent internal model (given **r**, what do we know about **x** or p_b_(**x**|**I**)?), or to apply the logic of Bayesian Encoding to task-specific quantities (construct **r** to encode a desired p(*s*|**I**)), but such examples are rare. In the remainder of this paper, we will associate Bayesian Encoding with task-independent internal generative models and Bayesian Decoding with variables in a task, and in the Discussion we will return to the possibility that task variables are explicitly represented as part of the brain’s internal model.

#### 2.3.2 Differing notions of likelihood

Another difference in philosophy is evidenced by divergent usage of the term “likelihood” (Figure 3). In the typical Bayesian Encoding setting, the term “likelihood” is reserved for the abstract relationship between internal model variables and sensory observations. For instance, one could speak of the “likelihood that this configuration of variables in the brain’s model generated the observed image,” or p_b_(**I**|**x**). This usage supports the idea that the quantity being computed is a posterior *over variables in a generative model of sensory data.* In the typical Bayesian Decoding setting, on the other hand, the “likelihood” refers to a relationship between stimuli and neural responses, p(**r**|*s*). This usage supports the idea that the quantity of interest is the posterior *over external stimuli in a task.*

#### 2.3.3 Differences in the relationship between distributions and neural activity

Bayesian Encoding models require two distinct assumptions: first, what is the source of the reference distribution to be encoded (e.g. what is the brain’s internal model p_b_(**x, I**)); and second, what is the linking hypothesis that maps probability distributions to neural activity (Figure 1)? This approach of starting with the encoded distribution abstracts away from the details of neural circuits that must actually *implement* inference. Take the model of Orbán et al. (2016) for example. In this work, the authors assume that neurons in primary visual cortex implement a sampling algorithm to encode the posterior distribution over latent variables in a Gaussian Scale Mixture model. This specifies the reference distribution. It is then assumed that, by some *unspecified* mechanism, the trajectory of a set of neurons’ membrane potentials over time traces out real-valued samples from the posterior, and that these membrane potentials elicit spikes through a nonlinear accumulation process. This specifies the linking hypothesis, or the map from the reference distribution to neural data. This model successfully reproduced a diverse set of known properties about V1 (Orbán et al., 2016), but it is not a mechanistic model. From a modeling standpoint, the way that an input image elicits neural activity is *mediated* by the reference posterior: an example of “downward causation” (Campbell, 1974; Yablo, 1992).

While for Encoding models there is a clear separation of computational model and neural link, they still of course beg the question of how inference in the computational model is implemented in neural circuits. Prior work has investigated the question of how biologically-plausible recurrent circuits could implement sampling (Bill et al., 2015; Probst et al., 2015; Aitchison and Lengyel, 2016; Petrovici et al., 2016; Dold et al., 2019; Echeveste et al., 2020) or message-passing (George and Hawkins, 2009; Beck et al., 2012; Raju and Pitkow, 2016; Grabska-Barwinska et al., 2013; Grabska-Barwińska et al., 2017; George et al., 2018) through their dynamics. However, in these examples there is typically a cost to increased biological plausibility, either by degrading the quality of the encoded distribution, or by degrading the match to empirical neural data.

Bayesian Decoding models, in contrast, do not distinguish between the uncertainty in an underlying probabilistic model and the uncertainty of a downstream brain area applying a Bayesian decoder to some neural activity. As a result, Decoding models replace the assumption about the link to neural activity with a *constraint* on the relationship between stimuli (*s*) and neural activity (**r**).

To illustrate this point, let us revisit one of the motivating examples for distributional codes of Zemel et al. and contrast the Encoding and Decoding approaches. Consider a rat who is placed into a water maze and must navigate to a hidden platform (Morris, 1984). Initially, the rat may be uncertain about which direction it is facing, e.g. if opposite walls of the maze look the same its correct belief about direction will be bimodal. Similar to the orientation of a grating, head direction is a scalar variable in [0, 2π] that we will call *s*. In the Encoding approach, one might begin by asking what is p_b_(*s*|**I**) according to an internal model of the environment, where **I** stands for the sensory cues the rat uses to orient itself. The distribution p_b_(*s*|**I**) determines how uncertain the rat *ought* to be, according to the internal model. Continuing the Encoding approach, one would then adopt a linking hypothesis (sampling, parametric, etc.) whereby p_b_(*s*|**I**) is encoded in neural activity **r**. In an Encoding model, the encoding of a distribution may be imperfect and lossy, or it may contain more information about the distribution than is being used by downstream circuits. In either case, the way a downstream circuit *uses* the neural activity will generally differ from a Bayesian decoder.

Applying the Bayesian Decoding framework to the same problem, we would say that the uncertainty in p(*s*|**r**) is the primary kind of uncertainty we should be concerned with, and that there is no distinction between this and the rat’s internal model. Crucially, this does not trivialize representations of uncertainty as “just” a matter of optimal decoding. In the Decoding approach, there may still be an ideal uncertainty that the rat *ought* to have when it is first placed into the maze; however the assumption is that this uncertainty is realized through the way **r** is tuned to its inputs **I**. That is, it is left to the brain (evolution, learning) to have carefully constructed tuning functions p(**r**|**I**), such that p(*s*|**r**) is equal to p_b_(*s*|**I**) (Ma et al., 2006). One way that the Encoding and Decoding perspectives can become identical, then, is when the decoded distribution p(*s*|**r**) equals the reference distribution p_b_(*s*|**I**). From the Encoding point of view, this requires that the encoding of p_b_(*s*|**I**) into **r** is lossless (or “efficient” in the terminology of Beck et al. (2012)). From the Decoding point of view, they are identical by assumption.

Finally, the preceding discussion points to an important practical difference between Encoding and Decoding philosophies in terms of how neural responses are interpreted by downstream areas. In a Decoding model, a downstream area implicitly applies Bayes’ rule to the neural responses arriving from an upstream area to extract information about a stimulus. In an Encoding model, on the other hand, upstream neural activity represents samples or parameters that are then processed by the downstream area according to an underlying approximate inference algorithm, which generally will *not* apply Bayes’ rule to the incoming activity directly. To put it another way, if one assumes that upstream neural activity encodes samples or parameters in an approximate inference algorithm, then there is an important conceptual difference between a downstream area that interprets upstream activity *as samples* or *as parameters* (as in Encoding models), and a downstream area that *decodes* the activity it receives by applying Bayes’ rule to the neural activity.

#### 2.3.4 Differing Empirical and Theoretical Motivations

Finally, distinguishing Bayesian Encoding and Bayesian Decoding allows one to be more precise on what data and what normative arguments motivate different theories. Bayesian Decoding can be motivated by the fact that humans and other species are empirically sensitive to uncertainty and prior experience, as in the classic psychophysics results on multi-modal cue combination (Ernst and Banks, 2002; Knill and Pouget, 2004; Alais and Burr, 2004; Körding, 2007; Angelaki et al., 2009; Pouget et al., 2013). The large literature on optimal or near-optimal Bayesian perception in controlled tasks motivates the question of how neural circuits facilitate Bayesian computations *with respect to stimuli in a task*, which are often scalar or low-dimensional. With the additional assumption that the neural representation of task-relevant aspects of stimuli is formatted to be easily decoded (e.g. linear and invariant to nuisance (Ma et al., 2006)), this line of reasoning has given rise to predictions for neural data. These predictions have since been largely confirmed for the representation of self-motion in dorsal medial superior temporal area (MSTd) (Fetsch et al., 2011, 2013; Hou et al., 2019). Bayesian Decoding is further motivated by experimental data showing a correspondence between non-parametric likelihood functions, neural noise, and behavioral indications of uncertainty (Walker et al., 2019).

Importantly, none of these results constitute direct evidence for inference with respect to an (usually high-dimensional) internal model of natural stimuli, as hypothesized in typical Bayesian Encoding theories (Rahnev, 2019; Koblinger et al., 2021). The three lines of support for Bayesian Encoding models are largely independent of the above motivations for Bayesian Decoding. First, Bayesian Encoding can be motivated by the purely normative argument that any rational agent that faces uncertainty *ought to* compute probability distributions over unobserved variables, as long as those variables directly enter into calculations of expected utility (Jaynes, 2003). Second, there is some empirical evidence for a key prediction of Bayesian encoding models: a general constraint on *all* well-calibrated statistical models is that the prior must equal the average posterior (Dayan and Abbott, 2001). Existing observations suggest that this constraint is satisfied in early visual cortex, as evidenced by changes in neural responses in primary visual cortex over development (Berkes et al., 2011) and task-learning (Haefner et al., 2016; Lange and Haefner, 2022). Third, there is empirical evidence for signatures of particular inference algorithms and particular internal models fit to natural stimuli. This approach has been employed by a series of sampling-based inference models and has successfully reproduced a wide range of neural response properties in early visual cortex (Orbán et al., 2016; Aitchison and Lengyel, 2016; Echeveste et al., 2020). A similar approach has also been taken by parametric models, where neural circuits have been hypothesized to implement the dynamics of a variational inference algorithm (Friston, 2005; George and Hawkins, 2009; Beck et al., 2012; Grabska-Barwinska et al., 2013; Raju and Pitkow, 2016; George et al., 2018; Lavin et al., 2018). We emphasize again that existing evidence for Bayesian-like behavior in psychophysical tasks only constitutes weak evidence in support of the idea that the brain computes distributions over variables in a task-independent internal model, as usually studied in the Bayesian Encoding literature (Rahnev, 2019; Koblinger et al., 2021).

### 2.4 Classification of existing models

Historically, sampling-based neural models have taken the Bayesian Encoding approach, asking how neurons could sample from the posterior distribution over variables in an internal model, while PPCs have primarily been studied in the context of inference of low-dimensional task-relevant quantities. However, this does not reflect a fundamental distinction between the two types of distributional codes. Parametric codes can and have been used in Bayesian Encoding models to approximate the posterior over variables in a generative model, including Probabilistic Population Codes (PPCs) (Beck et al., 2012; Grabska-Barwinska et al., 2013; Raju and Pitkow, 2016), Distributed Distributional Codes (DDCs) (Vertes and Sahani, 2018), and others (Friston, 2005; George and Hawkins, 2009; Lavin et al., 2018; George et al., 2018). Markov Chain Monte Carlo (MCMC) sampling has been used to explain perceptual bistability (Moreno-Bote et al., 2011; Gershman et al., 2012), which could be seen as a form of sampling-based Bayesian Decoding (cf. Hohwy et al. (2008)). To summarize, Table 1 provides a list of examples in each of the four categories defined by the sampling versus parametric and the encoding versus decoding axes. The fact that there is previous work in all four quadrants emphasizes that these are complementary distinctions.

### 2.5 Case Study: primary visual cortex (V1)

We now focus on primary visual cortex (V1) to provide a concrete example illustrating and further elaborating on our general points above. Focusing on area V1 has the advantage that much neurophysiological data exists, and both encoding and decoding approaches have enjoyed some success. We will first briefly describe existing work from both perspectives, and then use a simple example to show how they can lead to very different conclusions about the neural code. To that end, we will assume a Bayesian Encoding model that encodes the posterior over internal variables by sampling and show analytically how to derive the corresponding Bayesian Decoding model, obtaining a parametric representation (PPC) (Shivkumar et al., 2018).

#### 2.5.1 Bayesian Encoding models for V1

The starting point for the Bayesian Encoding approach, applied to V1, is an assumption about the brain’s generative model p_b_(**x, I**). That is, we must specify what is **x**, the variable assumed to be inferred and represented by V1 neurons, and how **x** is related to the sensory observations, **I**. For simple cells in area V1, Olshausen and Field proposed a linear Gaussian likelihood 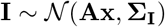 with a sparse independent prior p_b_(**x**) as the brain’s internal model Olshausen and Field (1996, 1997). (We use the notation 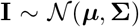 to indicate a random variable drawn from a multivariate normal distribution, and 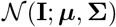 to denote its density function). In this model, the observed retinal image, **I**, is assumed to be a linear combination of “projective fields” (**PF***_i_*) plus unexplained pixel noise **Σ_I_**; the matrix **A** is a feature dictionary with projective fields as its columns: **A** = (**PF**_1_,…, **PF***_n_*). Each of the *n* projective fields is weighted by a single latent variable, **x** = (*x*_i_,…, *x_n_*)^⊤^. Intuitively, in this model, V1 activity is assumed to somehow represent beliefs about what values for **x** best explain a given retinal image, **I**.

The next assumption in the Encoding framework is about the neural code, or how the posterior distribution, p_b_(**x**|**I**), is represented by neural responses, **r**. Continuing the previous example, Olshausen and Field assumed that each *x_i_* was represented by a single neuron whose firing rate was proportional to the most probable value for *x_i_* given an image (maximum a posteriori, MAP): *r_i_* ∝ argmax*_x_i__* p_b_(**x**|**I**). In this model, a single neuron represents the most likely intensity with which a visual feature is present in the image. This is not a fully Bayesian Encoding model in the sense that only the MAP, but not the full posterior distribution p_b_(**x**|**I**) is encoded in neural responses. Empirical support for this model is based on the observation that learning (fitting) this model on natural images yields visual features (**PF***_i_*) that are localized, oriented, and band-pass filtered, implying neural responses and receptive fields with similar properties – just as observed empirically (Olshausen and Field, 1996).

Subsequent work has both modified and extended this generative model, and combined it with different neural codes. Hoyer and Hyvärinen proposed that neural responses can be understood as samples from the posterior in the same generative model to qualitatively explain the variability and mean-variance relationship of neural responses. Schwartz and Simoncelli extended the generative model to a Gaussian scale mixture model to explain the empirically observed contrast normalization of V1 responses, and Orbán et al. (2016) found agreement between the predictions of a Gaussian scale mixture model combined with neural sampling and a wide range of observations related to the stimulus-dependence of neural variability. Bornschein et al. (2013) proposed a variation of the generative model of Olshausen and Field using a nonlinear Gaussian likelihood and/or binary as opposed to continuous latents **x**, and Coen-Cagli et al. (2015) found that a further extension to the generative model in the form of a Mixture of Gaussian Scale Mixture model could explain center-surround interactions in V1. Finally, Haefner et al. (2016) combined the generative model of Olshausen and Field with the ideal observer model of a discrimination task to explain choice probabilities and task-dependent noise correlations of V1 neurons.

The key shared element of all these models is an explicit assumption about the computational variable **x** that is being represented, and how this variable is related to the sensory observations **I**. This model being adapted to natural inputs is an important constraint, and the model parameters are usually obtained by fitting the model to sets of natural images. These models are then general purpose and can be queried using natural inputs or images presented in a task. The fundamental framing is how V1 neurons *encode* the posterior over **x** given an arbitrary input, **I**.

#### 2.5.2 Bayesian Decoding models for V1

The starting point for the Bayesian Decoding approach, applied to V1, is a measurement of the conditional probability (or likelihood of s), p(**r**|*s*), for some stimulus *s* to which V1 neurons are tuned, and that is hypothesized to be represented, such as orientation. Importantly, this means that the measured likelihood is to some extent under experimental control, since the experimenter chooses what images correspond to each value of *s* (e.g. the size, contrast, or spatial frequency of a grating). In general, for an arbitrary *s*, this likelihood will be very complicated reflecting the fact that *s* cannot easily be decoded from **r** (e.g. object identity from V1). However, for V1 responses it has empirically been found that the optimal Bayesian decoder for orientation is approximately linear in spike counts and invariant to contrast, a classic nuisance variable (Graf et al., 2011). This finding has been interpreted as meaning that V1 activity “represents” orientation. In conjunction with the Poisson-like neural response variability in V1, this implies that the beliefs of a Bayesian decoder of orientation applied to the neural responses are part of the exponential family. Furthermore, the sufficient statistics are linear in the neural responses. Such a neural representation of a belief has been called a Probabilistic Population Code (PPCs) as introduced by Ma et al..

The same logic applies to other candidates for *s* that modulate the responses of V1 neurons in a straight-forward manner, such as spatial frequency or location. The key element of the Bayesian Decoding approach is taking the perspective of downstream circuits trying to extract information about *s* from V1 activity: how is the information about *s* formatted in V1 activity, and is p(*s*|**r**) “simple”? In contrast to the Bayesian Encoding perspective which justifies its choice of **x** by its fit to natural images and its ability to predict neural responses, the Bayesian Decoding perspective justifies its choice of *s* by desirable properties of an efficient decoder, e.g. linearity and invariance to nuisance variables.

#### 2.5.3 Example model where decoding a stimulus *s* from encoded samples results in a PPC

Since our main points are conceptual in nature, we will develop the link between the Encoding and the Decoding approach for the simple case of a linear Gaussian model with a Gaussian prior, under the assumption of a sampling-based neural code. These simplifying assumptions make the difference between Encoding and Decoding clear and analytically tractable, but are not meant to maximize biological plausibility. For instance, the posterior variance in this model is independent of **I**, whereas it is well-known that neural response variance is stimulus-dependent, and this effect is captured by neural sampling models with less trivial generative models (Orbán et al., 2016; Bányai et al., 2019; Festa et al., 2019). Beginning with a more complicated Encoding model would lead to a more complicated relationship to Decoding models (e.g. where the Decoder is more complex, e.g. a nonlinear PPC, or not even in the exponential family). Importantly, the core of our argument remains: that an Encoding model based on one type of neural code (e.g. sampling) and a Decoding model based on another type (e.g. parametric) need not be in contradiction with each other, and offer complementary perspectives on the same system.

Given an image, **I**, we assume that V1 neurons *encode* the posterior p_b_(**x**|**I**) by sampling *t* values from from the posterior distribution, **x**^(*t*)^ ~ p_b_(**x**|**I**) ∝ p_b_(**I**|**x**)p_b_(**x**) where p_b_(**x**) is the brain’s prior over **x** (Hoyer and Hyvärinen, 2003). We assume that responses from a population of n neurons correspond to samples from the posterior over **x**, so that at each instant, the population response, **r**^(*t*)^, equals the sample **x**^(*t*)^. Each sample of *x_i_* (or *r_i_*) represents the brain’s instantaneous belief about the intensity of the feature **PF***_i_* in the image.

We will now apply the Bayesian Decoding approach to the sequence of samples produced by the sampling-based Encoding model described above. An ideal observer applies Bayes’ rule to infer p(*s*|**r**^(1)^,…, **r**^(*t*)^) using knowledge of the probabilistic relationship between samples (**x** or **r**) and *s*:

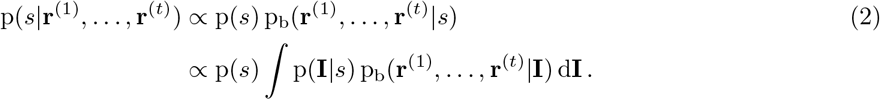

That is, the optimal decoder combines knowledge of (i) how likely an image **I** is to generate a set of samples of **x** (or **r**), and (ii) how likely a stimulus value *s* is to generate an image **I**. In general, this decoded distribution over *s* may be arbitrarily complex and intractable. One factor that is under experimental control is the”template” function **T**(*s*) which renders an image, such as a grating with orientation *s*. This provides the link between *s* and **I** in equation (2). In our model, we assume that the input the brain receives is a noisy version of that template (Figure 4).

**Figure 4:**
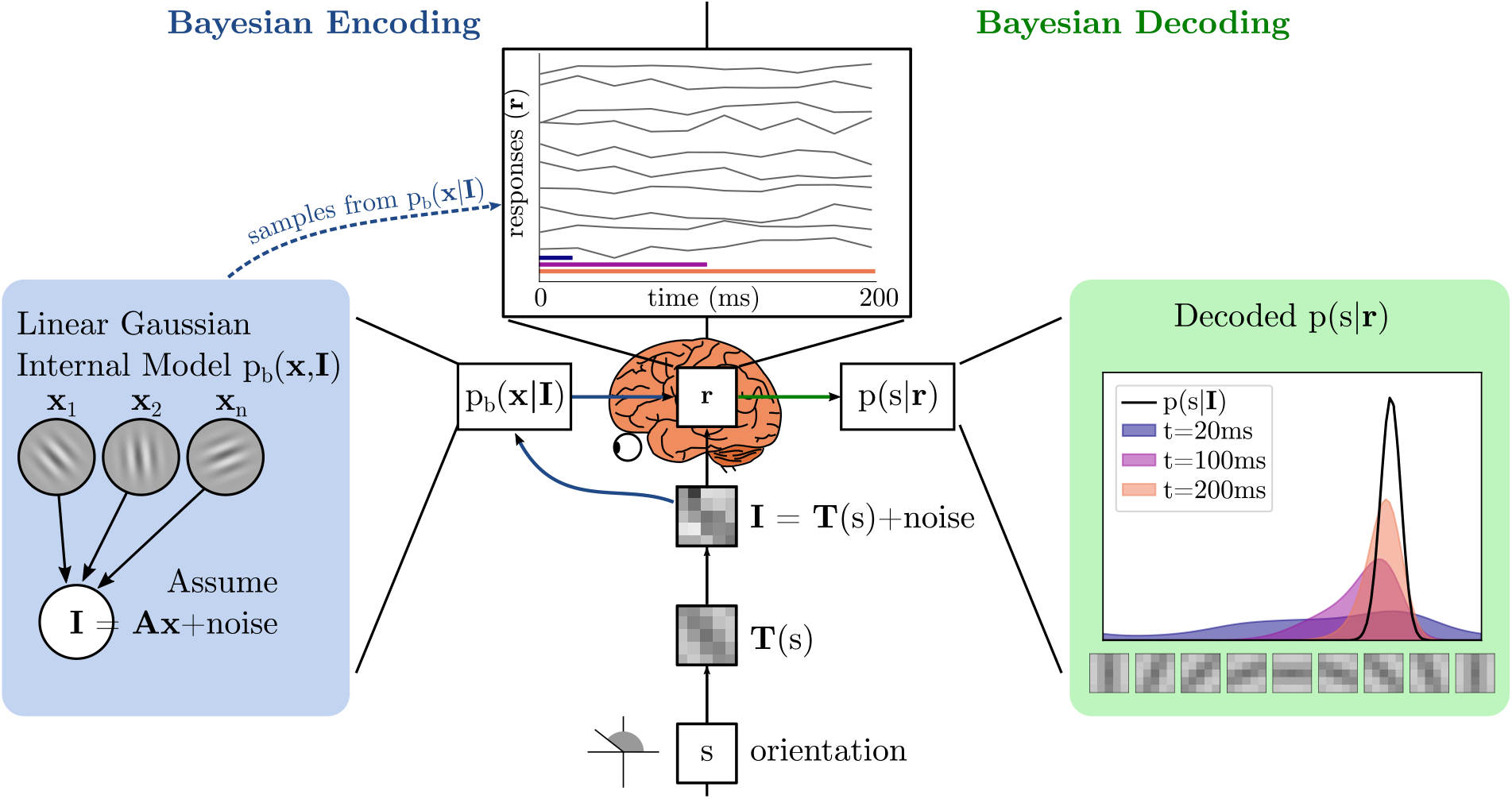
Encoding by sampling followed by decoding of orientation from the samples in a simplified model. As in Figure 3, Encoding elements are on the left, and Decoding elements are on the right. In our example model, the brain performs sampling-based inference over **x** in a probabilistic model of images, here a Linear Gaussian model. In a given experiment, the image is generated according to an experimenter-defined process that turns a scalar stimulus *s*, e.g. orientation, into an image observed by the brain. To simplify, neural responses **r** are assumed to reflect instantaneous real-valued samples of **x** drawn from the posterior p_b_(**x**|**I**). In our simulation we drew 10 samples and assumed 20ms per sample. The samples drawn from the model are then probabilistically “decoded” to a probability distribution over *s*. This distribution sharpens as more samples are observed. The optimal decoder for any *t* is a linear PPC.

The first simplification to the general form of the optimal decoder in (2) we can derive, under the assumption of a Gaussian likelihood, is to show that the posterior over *s* depends only on the mean rate of **r** (i.e. a rate code rather than temporal code):

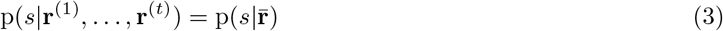

where 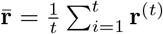 is the mean response after *t* samples (Supplemental Text). Any decoder that obeys (3) can be seen as a kind of *parametric code* over *s*, where the rates 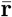 are the parameters. A second convenient property for a decoder to have is if the optimal decoder is in an exponential family, or

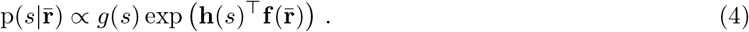

Whenever (4) is true, then we would say that the neural activity forms a particular kind of parametric code called a *Nonlinear PPC* over *s*. A final convenient property for a decoder to have is if 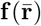 is linear:

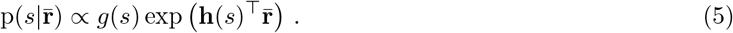

This is the definition of a *Linear PPC* over *s* (Ma et al., 2006)^1^.

In our simplified Encoding model, we can analytically derive the optimal decoder of the experimenter-defined *s* conditioned on neural responses, where those neural responses are generated by sampling from the Linear Gaussian internal model described above (derivation in the Supplemental Text). We find that the optimal decoder is, in fact, a Linear PPC over *s* as defined in (5)!^2^ This sequence of steps from equation (2) through (5) suggests a general way to derive the Bayesian Decoding model implied by a given Bayesian Encoding model.

As we discussed earlier, Encoding models typically (but not necessarily) consider inference in a task-independent internal model, while Decoding models typically (but not necessarily) consider inference of low-dimensional task-specific quantities. The model we described in this section is “typical” in this sense: inference of **x** is constructed to be task-independent, while the decoder of *s* given **r** depends inextricably on the “template function” **T**(*s*), which is under the control of the experimenter. The kernels **h**(*s*) will be different for gratings of different size and spatial frequency, for plaids, or for different objects. This example shows how a Bayesian Decoding model for *s*, implied by a task-independent Bayesian Encoding model, can nonetheless be *experiment-dependent*. This points to a potentially empirically decidable question: do downstream areas such as V2 interpret V1 activity like a Bayesian Decoder, or do they interpret V1 activity as representing a posterior over a set of latents, **x**?

## 3 Discussion

We have identified and characterized a previously unstated difference between approaches to constructing Bayesian neural models: Bayesian Encoding and Bayesian Decoding. This distinction is orthogonal to existing and much-debated distinctions like whether neural responses reflect parameters or samples of the inferred distribution. Making the distinction between Bayesian Encoding and Bayesian Decoding explicit provides new insights into the long-standing debate about the nature of the neural code. Importantly, we have demonstrated that these two approaches can give rise to different but compatible models of the same neural circuit, underlining the point that Bayesian Encoding and Decoding models are complementary, and not mutually exclusive. The complementary nature of these approaches has direct implications for both theoretical debates and the correct interpretation of empirical data.

Our example model sheds light on the much-debated question of whether neural responses are more closely related to parameters of the encoded probability distribution, as in probabilistic population codes (PPC; Ma et al. (2006)) and in distributed distributional codes (DDC; Vertes and Sahani (2018)), or to samples from the distribution as in neural sampling (reviewed in Fiser et al. (2010); Pouget et al. (2013); Sanborn (2015); Gershman and Beck (2016)). In our example, the Bayesian Decoding model implies a (parametric) PPC while, by construction, neural responses in the Bayesian Encoding model represent samples, demonstrating that it is possible that the very same neural responses are compatible with both depending on perspective.

Our model is a constructive proof that Encoding and Decoding models *can* be compatible on the same data, but this is will not be true in general. For instance, non-Gaussian Encoding models will not generally form a linear PPC from the decoding perspective, or only for specific sets of stimuli, or they may be decodable only as a nonlinear PPC. Generalizing from our specific example, the key question is, which families of Encoding models, consisting of both p_b_(**I, x**) and an assumption about the link to neural responses, are compatible with which families of Decoding models, consisting of p(*s*|**r**) and p(**I**|*s*)? These will come in pairs – each family of Encoding models defines a family of compatible Decoding models, and vice versa. Identifying these pairs of compatible model families is a theoretical question with important implications for the interpretation of empirical data: while Encoding and Decoding models traditionally have been supported (and falsified) by different kinds of empirical data (see section 2.3.4), understanding their link will allow us to bridge that divide. For instance, if an Encoding model implies a particular family of Decoding models, then data that falsifies that Decoding family will also falsify the Encoding family. Similarly, if a Decoding model is only compatible with a family of Encoding models that is too constrained to effectively model natural inputs, then that would pose a challenge for the Decoding model. As an example of this kind of argument, Orbán et al. note in supplemental analyses that their sampling model appears to be empirically consistent with a contrast-invariant linear PPC over orientation, but that “linear decoding of population responses will significantly fall short of being optimal” once more complex tasks are considered (Orbán et al., 2016).

More generally, our arguments also raise questions about what makes a neural code “distributional”, i.e. representing a whole distribution rather than just a point estimate, and what would constitute empirical evidence for it. While the Bayesian Encoding model in our example assumed a sampling-based representation of the posterior over **x**, consider a reduced, non-distributional version in which neural responses are proportional to a point estimate of **x** such as its mean or mode (Olshausen and Field, 1996). This would be a poor Bayesian Encoding model in the sense that the full p_b_(**x**|**I**) distribution is not recoverable from **r**. Yet, even this reduced model gives rise to a probabilistic code (PPC) over *s*. Such a *point estimate* code over variables in the brain’s internal model would still enable many of the apparently Bayesian behaviors observed in low-dimensional psychophysics tasks and used to motivate Bayesian Decoding theories, as discussed in section 2.3.4 above. Another example of non-Bayesian encoding but Bayesian decoding is given by Orhan and Ma. It would therefore be a mistake to treat empirical evidence for near-optimal or near-Bayesian behavior in a particular task alone as evidence that the brain represents probability distributions over variables in an internal model of sensory inputs (Rahnev, 2019; Koblinger et al., 2021). The distinction between Bayesian Encoding and Bayesian Decoding might thus productively add to the open philosophical question: “if perception is probabilistic, why does it not seem probabilistic?” (Block, 2018; Rahnev et al., 2020).

The seminal paper by Zemel et al. (1998) introduced the concept of encoding (and decoding) general probability distributions in (and from) neural activity. Most work over the following 20+ years typically focused on either Encoding or Decoding, as shown by Table 1, despite Zemel et al. considering both perspectives as tightly linked. This divergence was likely strengthened by the fact that Encoding studies almost exclusively considered internal latent variables (**x**), while work taking the Decoding perspective considered distributions over task-defined variables (*s*). From today’s perspective, the encoding formalism of Zemel et al. and its application in Sahani and Dayan (2003) maps naturally onto our Bayesian Encoding category. Furthermore, it is philosophically closely aligned with the other studies in this category, and shares with them the idea that implied decoders that are non-Bayesian (but note that “decoding is only an implicit operation that the system need never actually perform” Zemel et al. (1998)). Interestingly, while Zemel et al. (1998) discounted the possibility of optimally decoding the encoded distribution using Bayes’ rule as too inflexible, almost all later studies that took the decoding approach were based on Bayes’ rule, and now form our Bayesian Decoding category.

The key step in our example system above which allowed us to interpret samples of **x** as a PPC was to construct the PPC over a different variable: *s*. This raises the question: what if *s* is part of of the brain’s internal model? One possibility is that “orientation” (or any other *s* in a task) is a useful abstraction of natural stimuli, in which case it may have been learned (or evolved) and may permanently be a part of the brain’s internal model. Another possibility is that orientation (or any other s) is part of the brain’s internal model because the brain changes its internal model as the result of learning the present task (Haefner et al., 2016; Lange and Haefner, 2022). Echoing section 2.3.3 above, even if *s* is part of the brain’s internal model, Bayesian Encoding and Decoding models would nonetheless differ in their approach to the question of how neural responses, **r**, relate to the distribution on *s*. Bayesian Encoding models would begin with a generative model of sensory input **I** from *s* (and possible other internal variables **x**) and ask how the true posterior p_b_(s, **x**|**I**) is represented by neural responses **r**. Bayesian Decoding models, on the other hand, would investigate the relationship between *s* in the world and evoked neural responses, p(**r**|*s*), and study a different kind of posterior, p(*s*|**r**), which takes the perspective of the experimenter, or possibly the rest of the brain trying to read out *s* from **r**. If the *decoded* distribution, p(*s*|**r**), matches the ideal *encoded* distribution, p_b_(*s*|**I**), then the code for *s* is said to be *efficient* (Beck et al., 2012).

The choice of variable which is assumed to be inferred, also impacts the interpretation of neural variability. In our example above, neural variability is directly related to the uncertainty in the posterior over **x**. In contrast, the uncertainty over *s* encoded by the Bayesian decoding model is unrelated to the neural variability, depending on the samples only through their *mean,* rather than their *variance.* Given sufficiently many samples, the uncertainty over *s* is only determined by the noise in the channel between experimenter and brain (**Σ**_e-b_). This is an important point for experiments that seek to test the neural sampling hypothesis by relating neural variability and “uncertainty”: in our example model, only uncertainty over **x** but not over *s* manifests as neural variability, while *s* is the variable most commonly and naturally manipulated in an experiment.

The issues raised in this paper for models of visual perception also have implications for Bayesian models of cognition, where ideas related to sampling (Vul and Rich, 2010; Sanborn et al., 2010; Lieder et al., 2014; Vul et al., 2014; Sanborn and Chater, 2016; Lieder et al., 2017; Zhu et al., 2020), variational inference (Hohwy et al., 2008; Daw et al., 2008; Sanborn and Silva, 2013), or both (Lange et al., 2021) have been invoked to explain a wide variety of heuristics and biases (reviewed in Sanborn (2015); Griffiths et al. (2012b)). Here, too, it is important to distinguish between probabilistic models of the world that are posited to exist in a subject’s mind (as is typical in Bayesian Encoding) from experimenter-defined models of a particular task (as is typical in Bayesian Decoding). Closely related is the distinction drawn by Knill and Richards between the “inference problem” (what the brain infers in the internal model it assumes) and the “information problem” (what information is available in the world) (Knill and Richards, 1996, Chapter 1). For example, Vul et al. argue that certain deviations from Bayes-optimal behavior can be explained as the result of basing decisions on a single Monte-Carlo sample. However, it is conceivable that what appears to be a single point-estimate sample over a quantity relevant to a task may, in fact, be a local, perhaps unimodal distribution over a detailed internal model, as in variational approximations. It is further conceivable that multiple “samples” correspond to a mixture of variational approximations over an internal model (Jaakkola and Jordan, 1998; Lange et al., 2022). Conversely, a single high dimensional point estimate of an internal model may be sufficient to facilitate apparently Bayesian behavior with respect to a low-dimensional task. Changing our reference frame from internal models to experimenter-defined tasks may make samples appear as variational parameters, or vice versa.

Koblinger et al. (2021) recently posed the question whether uncertainty in the brain is represented “constitutively”, i.e. about many variables regardless of their relevance for a specific task, or “opportunistically,” only about task-relevant variables. While this distinction appears related to the difference between Bayesian Encoding and Decoding with respect to their task-independence, there are important differences. While Encoding and Decoding models have so far mostly been applied in task-independent and task-dependent contexts, respectively, to what degree representations of uncertainty are task-specific is an empirical question that can be productively asked within both the Encoding and Decoding approaches. For instance, Bayesian Encoding models of object recognition may differ in whether they propose that the brain represents posteriors only over task-relevant object identities, or all possible objects. One can similarly imagine both “constitutive” and “opportunistic” Decoding models. For instance, the toy example model we presented above is an opportunistic Decoding model, where *s* is determined by the experimenter, and **r** is only said to represent a distribution over *s* in that task context. In a constitutive Decoding model, the representation of a distribution about one quantity, like p(orientation| **r**), would potentially interact with representations of other quantities, like p(location| **r**), regardless of the immediate task-relevance of each.

Walker et al. (2022) pointed out a distinction between “descriptive” versus “process” approaches to the study of neural representations of uncertainty. In their classification, the “descriptive” approach derives estimates about the observer’s subjective uncertainty from either presented stimuli or recorded behavioral reports. The “process” approach, on the other hand, derives an estimate of uncertainty from neural responses. To what degree this classification is related to the Bayesian Encoding and Decoding approaches, respectively, is unclear, and likely depends on additional assumptions, e.g. about the relationship between behavioral reports and reference posterior in the Encoding approach, p_b_(**x**|**I**), and about the nature of the model used to infer uncertainty from neural responses.

To conclude, the Bayesian Brain Hypothesis is not a single idea, but a collection of computational models, philosophical ideas, and explanations for a variety of empirical data. It is a *framework* rather than a *theory* (Griffiths et al., 2012a). Bayesian Encoding and Bayesian Decoding are complementary approaches to constructing concrete models within the Bayesian Brain framework. These two approaches have been a major source of variation among models, and their complementary nature has previously gone unnoticed. We hope that these insights will lead to clearer and more productive discussions on the nature of inference in the brain, both in terms of neural representations of probability and in terms of behavior.

## Code availability

Two panels in Figure 4 were generated by simulation. The code is available at https://github.com/haefnerlab/bayesian-encoding-decoding.

## S Supplemental Text

### S.1 Derivation of decoded posterior for Gaussian prior over x

In this section we provide a brief derivation of the optimal posterior over an experimenter-defined *s* conditioned on samples of internal-model variables **x**, where the brain’s internal model p_b_ (**x, I**) is assumed to be a linear Gaussian model with a Gaussian prior over **x**. The use of a Gaussian prior over **x** is a further simplification of the derivation in Shivkumar et al. (2018). Formally, the setup is as follows:

1. Assume that the scalar *s* (such as orientation) gives rise to observed images **I** as

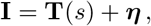

where **T**(*s*) is a “template” function (such as a grating image), and ***η*** is zero-mean Gaussian-distributed pixel noise with covariance **Σ**_e-b_.
2. Assume that the brain’s internal model, p_b_(**x, I**), factorizes as p_b_(**x**)p_b_(**I**|**x**), where the prior is Gaussian,

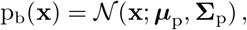

and images are assumed to be generated as a linear combination of basis vectors,

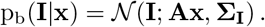
3. **Bayesian Encoding Model:** Assume that, conditioned on **I**, the brain samples {**x**^(*i*)^} ~ p_b_(**x**|**I**), and that each value of **x** corresponds to a neuron, so that **r**^(*i*)^ = **x**^(*i*)^. (We will use “**r**^(*i*)^” to denote the vector of neural activity at time *i*, and **r** = {**r**^(1)^,…, **r**^(*t*)^} to denote all neural activity in the relevant population up to time *t*.)
4. **Bayesian Decoding Model:** We will derive the Bayesian decoder of *s* given {**r**^(1)^,…, **r**^(*t*)^}.

We are interested in the optimal decoder of *s* after *t* time has elapsed, or p(*s*|**r**). By Bayes’ rule, this is proportional to p(**r**|*s*)p(*s*) = p(**x**^(1)^,…, **x**^(*t*)^|*s*)p(*s*). That is, the quantity we must compute in order to optimally decode p(*s*|**r**) is the probability of seeing a given set of samples, **x**^(1)^,…, **x**^(*t*)^, provided a value of *s*.

Since *s* affects **r** through **I**, this likelihood function can be evaluated by marginalizing across all possible images

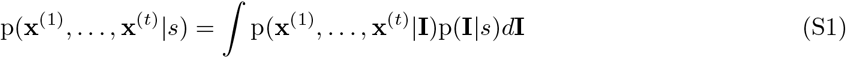

We know from our definition that

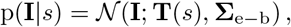

and the posterior probability of all *t* independent samples for a given **I** is

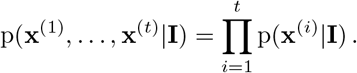

Under the simplifying assumption that both p_b_(**x**) and p_b_(**I|x**) are Gaussian, the brain’s internal model posterior, p_b_(**x**|**I**) is also Gaussian,

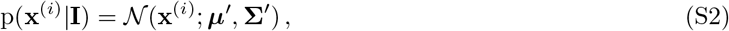

where

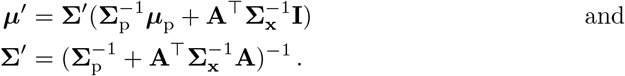

Note that the only dependence on **I** (and therefore on *s*) is through ***μ’***.

Equation (S2) gives the probability of seeing a single sample **x**^(i)^ given **I**. The probability of all *t* samples is

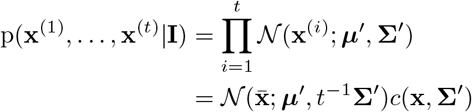

where 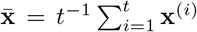 is the average of samples up to time *t*, and *c*(**x, Σ’**) is a term that depends on **x**^(1)^,…, **x**^(*t*)^ and on ***Σ’*** but not on ***μ’*** (and therefore not on **I**, so it can be dropped later).

We can now evaluate the integral in (S1) to get the probability of *t* samples for a given *s*:

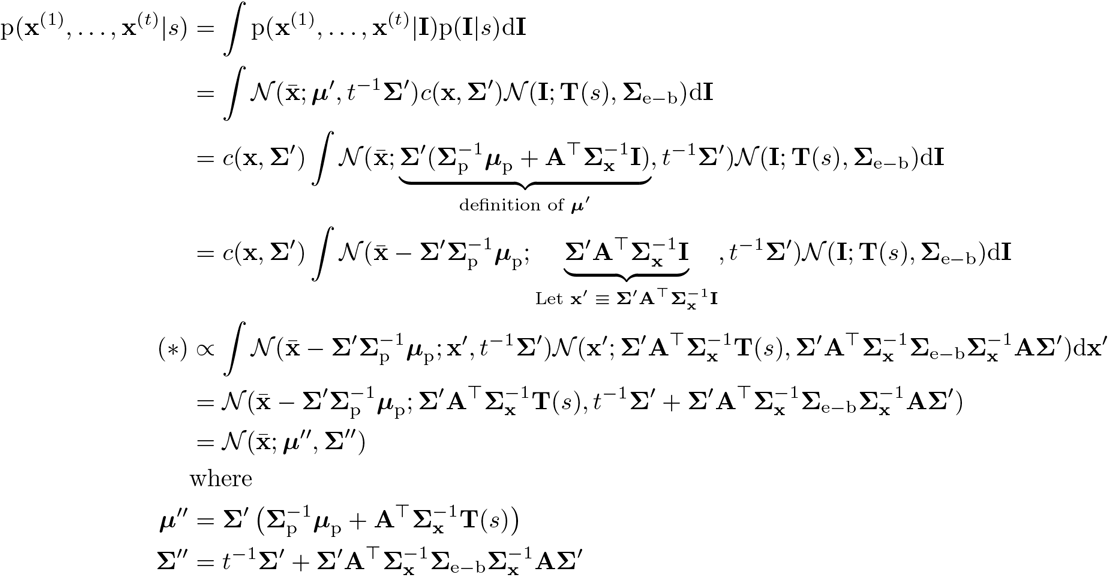

In the line marked (*), we changed variables to switch from an integral over **I** to an integral over **x**’. This line is a proportionality because we also dropped terms that do not depend on *s*, including the Jacobian term from the change of variables, since later we will use this expression as a likelihood function of *s*.

Expanding the definition of 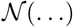, we can now write the posterior over *s* given **x**^(1)^,…, **x**^(*t*)^ as

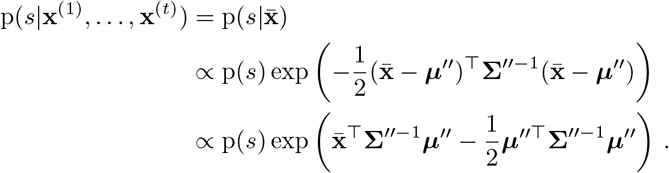

Substituting 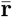 for 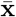 and rewriting in terms of a Linear PPC, this is

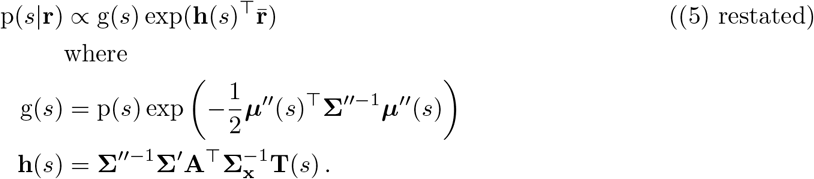

## Footnotes

1 PPCs also place restrictions on nuisance variables which we have omitted here.

2 Further discussion of the nature of this PPC and its relation to the parameters of the internal model can be found in Shivkumar et al. (2018).

